# Inserting Cre Recombinase into the Prolactin 8a2 gene for Decidua-Specific recombination in Mice

**DOI:** 10.1101/2022.01.28.478238

**Authors:** Mackenzie J. Dickson, Yeong Seok Oh, Artiom Gruzdev, Rong Li, Nuria Balaguer, Andrew M. Kelleher, Thomas E. Spencer, San-Pin Wu, Francesco J. DeMayo

## Abstract

An estimated 75% of unsuccessful pregnancies are due to implantation failure. Investigating the causes of implantation failure is difficult as decidualization and embryo implantation is a dynamic process. Here we describe a new decidua specific iCre recombinase mouse strain. Utilizing CRISPR/Cas9-based genome editing, a mouse strain was developed that expresses iCre recombinase under the control of the endogenous prolactin family 8, subfamily a, member 2 (Prl8a2) promoter. iCre recombinase activity was examined by crossing with mTmG or Sun1-GFP reporter alleles. iCre activity initiated reporter expression at gestational day 5.5 in the primary decidual zone and continued into mid-gestation (gestational day 9.5), with expression highly concentrated in the anti-mesometrial region. No reporter expression was observed in the ovary, oviduct, pituitary, or skeletal muscle, supporting the tissue specificity of the Prl8a2iCre in the primary decidual zone. This novel iCre line will be a valuable tool for in vivo genetic manipulation and lineage tracing to investigate functions of genetic networks and cellular dynamics associated with decidualization and infertility.

## Introduction

Implantation is a dynamic process that must occur during the window of receptivity for proper embryo development and pregnancy establishment. An estimated 30% of spontaneously conceived embryos are lost in the pre-implantation stage while more than 50% of in vitro derived embryos fail to implant (Teklenburg et al., 2010). In fact, an estimated 75% of failed pregnancies are due to implantation failure (Wilcox et al., 1988). Implantation failure represents a major limiting factor in successful pregnancy establishment.

An imperative step of implantation and pregnancy establishment is decidualization, which supports placental development and embryo growth. Stromal and epithelial cell crosstalk regulate transcriptomic changes and morphological remodeling of the endometrial stromal fibroblast cells into decidual secretory cells (Gellersen et al., 2007). This process of cell differentiation, decidualization and placenta establishment in the endometrium is vital for embryo survival and plays a role in trophoblast invasion, as well as protection against oxidative stress and maternal immunological rejection. Pregnancy and parturition are indicative of successful decidualization, however failure of proper decidualization and implantation, may lead to a multitude of problems including pre-eclampsia and preterm birth (Dimitriadis et al., 2020; Gellersen & Brosens, 2014; Ng et al., 2020).

Due to their transitory stage and critical value to embryo survival, studying decidual cells in vivo is challenging. For ethical concerns, it is not possible to study human endometrial decidualization in women and instead many studies utilize endometrial stromal cell cultures to study biological mechanisms. Investigators can isolate primary culture from uterine biopsies or use immortalized human endometrial stromal cells (Irwin et al., 1989; Krikun et al., 2004), however, in vitro conditions do not necessarily recapitulate what occurs in vivo. Rodent models are valuable as they also undergo decidualization for embryo implantation. While there is a mouse strain containing a Cre recombinase in the uterine stromal cells, *Amhr2*^*tm3(cre)Bhr*^, the Cre recombinase is also active in other cell types of the reproductive tract (Hernandez Gifford et al., 2009), which ultimately limits the interpretation of experimental results.

Specific key regulators of decidual cell signaling include members of the prolactin (PRL) superfamily (Handwerger et al., 1992; Jabbour & Critchley, 2001; Orwig et al., 1997; Telgmann & Gellersen, 1998). In humans and rodents, decidual cells express and produce PRL (Lee & Markoff, 1986), however in rodents there are an additional three members of the PRL superfamily expressed in uterine decidua including Prl family 8, subfamily a, member 2 (Prl8a2; also known as decidual Prl-related protein (dPRP)); Prl-like protein B (PLP-B), and PLP-J (Roby et al., 1993; Soares et al., 2007).

Our objective was to make a mouse strain with a decidua specific iCre (codon-improved Cre) recombinase. Utilizing CRISPR/Cas9-based genome editing, we created a strain that expresses iCre recombinase under the control of the endogenous *Prl8a2* promoter. Our results reveal that the *Prl8a2-iCre* allele is expressed in the uterine decidua following embryo attachment. Further, we demonstrate the tissue-specificity of this strain with reporter mice and histology.

## Results and Discussion

To create a new decidua-specific mouse strain Prl8a2iCre, CRISPR/Cas9-based genome editing was employed and the mouse codon-improved Cre recombinase (iCre) was inserted into the *Prl8a2* locus of embryonic stem (ES) cells. The iCre replaced a portion of the endogenous exon 1 and portion of intron 1 (**Figure 1**). Successfully targeted ES cells were subject to blastocyst injections to derive chimeric mice. Male chimeric mice were crossed with C57BL/6 female for germline transmission breeding to generate heterozygous mice that carry one copy of the *Prl8a2-iCre* allele. The resulting strain should express iCre recombinase under the control of the endogenous *Prl8a2* promoter.

**Figure 1.**
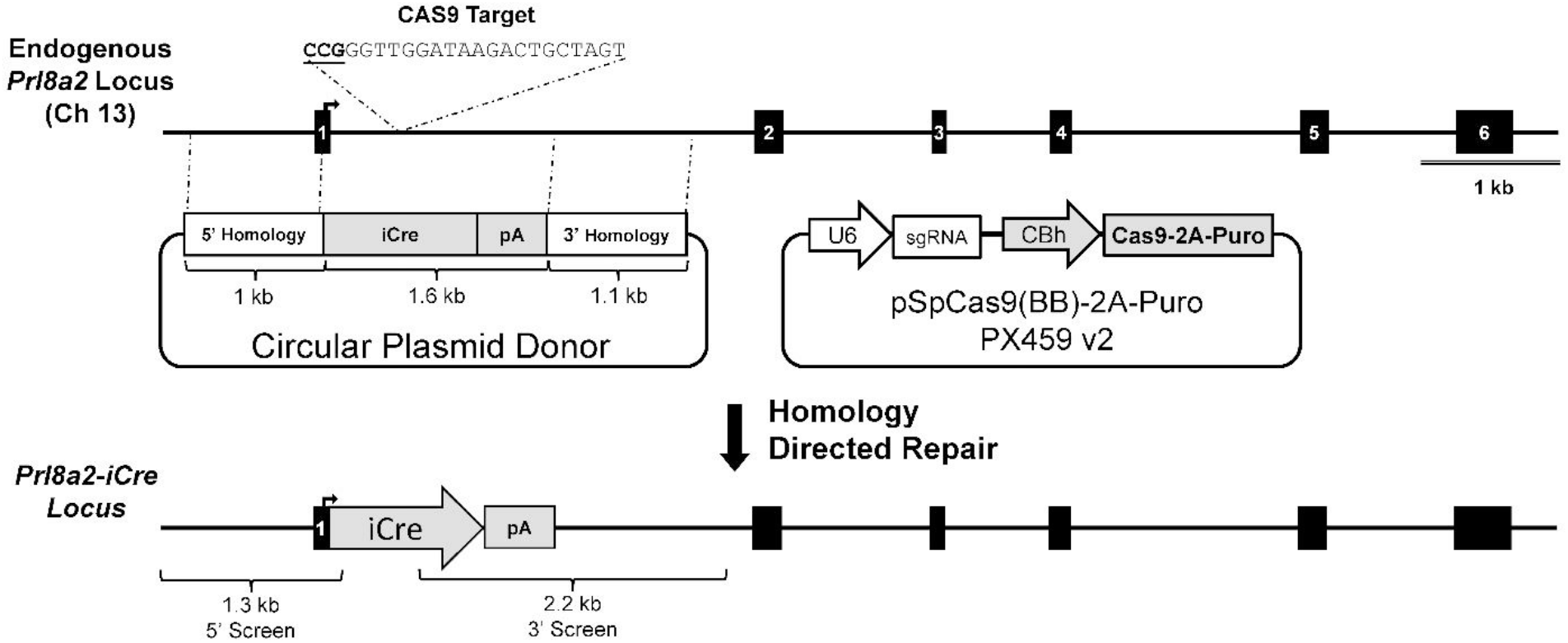
Targeting strategy to make the Prl8a2iCre allele. CRISPR/Cas9-mediated targeting strategy to generate the Prl8a2-iCre allele. The iCre ORF and polyA signal were inserted immediately downstream of the start codon in exon 1. The overall length of the locus was maintained by replacing 1.6 kb of endogenous sequence, mainly intron 1-2, with the 1.6 kb iCre-polyA genetic payload. The targeted locus was sequenced to confirm proper in-frame insertion of the iCRE ORF.

To examine the spatial and temporal distribution of cells expressing iCre driven by *Prl8a2* during early pregnancy, the *Prl8a2*^*i*Cre/+^ mice were mated with two different reporter strains; mTmG reporter mice (Muzumdar et al., 2007) and *Sun1*^LsL/+^ mice (Mo et al., 2015). Cells that were actively expressing or previously expressed the iCre recombinase constitutively manifested green fluorescence protein (GFP) signals in cytoplasm (mTmG) or nuclear membrane (*Sun1*^*LsL/+*^). Histological assays of uterine cross-sections from *Prl8a2*^*i*Cre/+^ mTmG mice showed iCre activity was detected via GFP expression in cytosol at GD 5.5 in the primary decidual zone (**Figure 2A**) into mid-gestation (**Figure 2B-C**). Of note, Prl8a2iCre activity can be induced on the anti-mesometrial side of the uterus five days after artificial decidualization, indicating that iCre activity and localization are independent of embryo-derived factors (**Figure 2D**). The use of a second reporter mouse line, *Sun1*^LsL/+^, to generate *Prl8a2*^*i*Cre/+^ *Sun1*^LsL/+^ mice verified iCre activity via GFP expression in the nuclear membrane of decidual cells in the anti-mesometrial region at gestational day 7.5 of pregnant mouse uterus (**Figure 3**).

**Figure 2.**
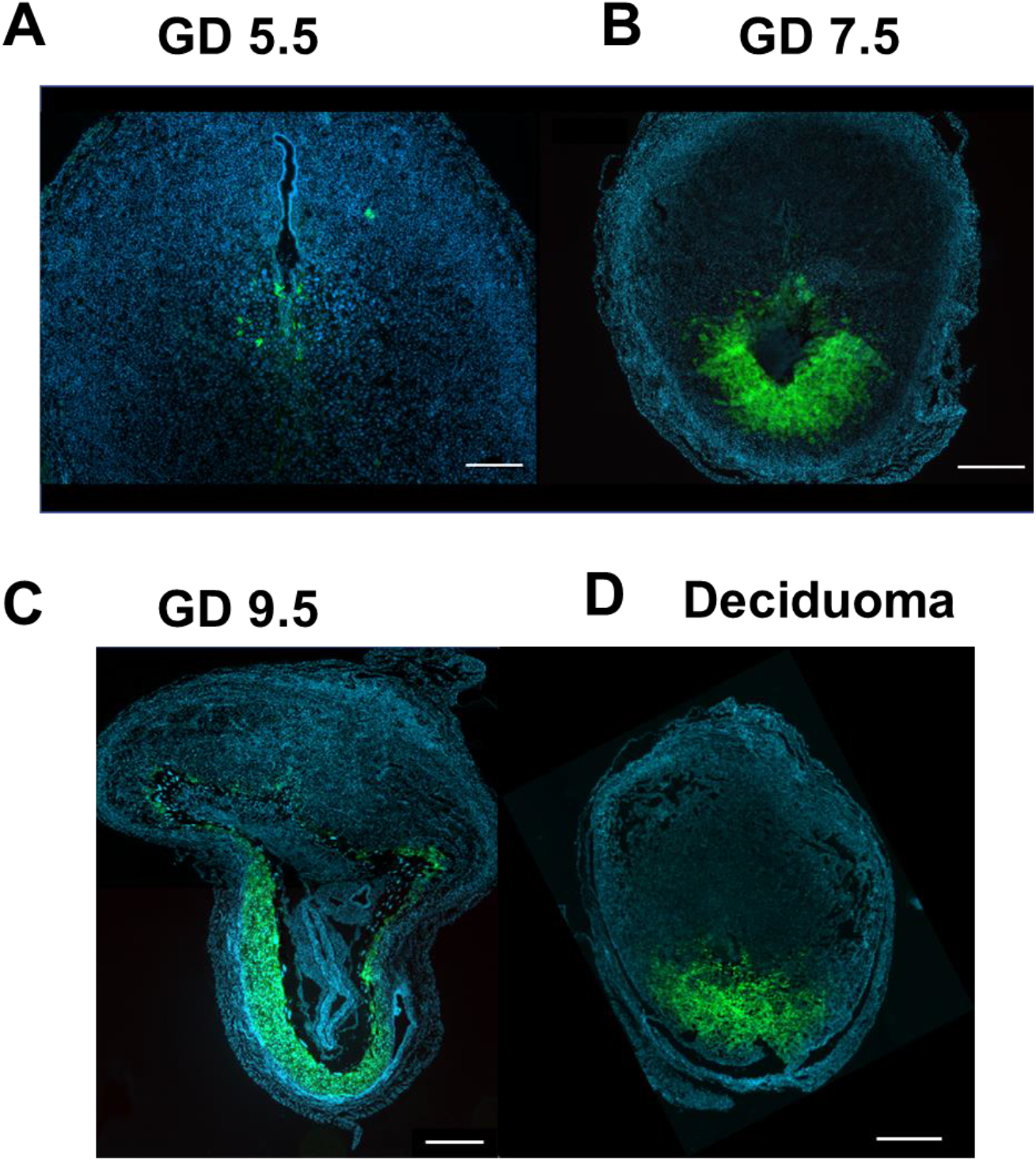
Time course of uteri from gestational days 5.5, 7.5, 9.5, and deciduoma in *Prl8a2*^*iC*re/+^ mTmG mice. *Prl8a2*^*i*Cre/+^ mTmG mice were mated with wild-type males and collected at various days of gestation to determine iCre activity in the uterus. *Prl8a2*^*iC*re/+^ activity is detectable at gestational day (GD) 5.5 (**A**) and increases at GD 7.5 (**B**). *Prl8a2*^*i*Cre/+^ is still apparent at GD 9.5 (**C**). *Prl8a2*^*i*Cre/+^ activity is present when artificial decidualization (**D**). Blue indicates DAPI nuclear stain and green is EGFP stain.

**Figure 3.**
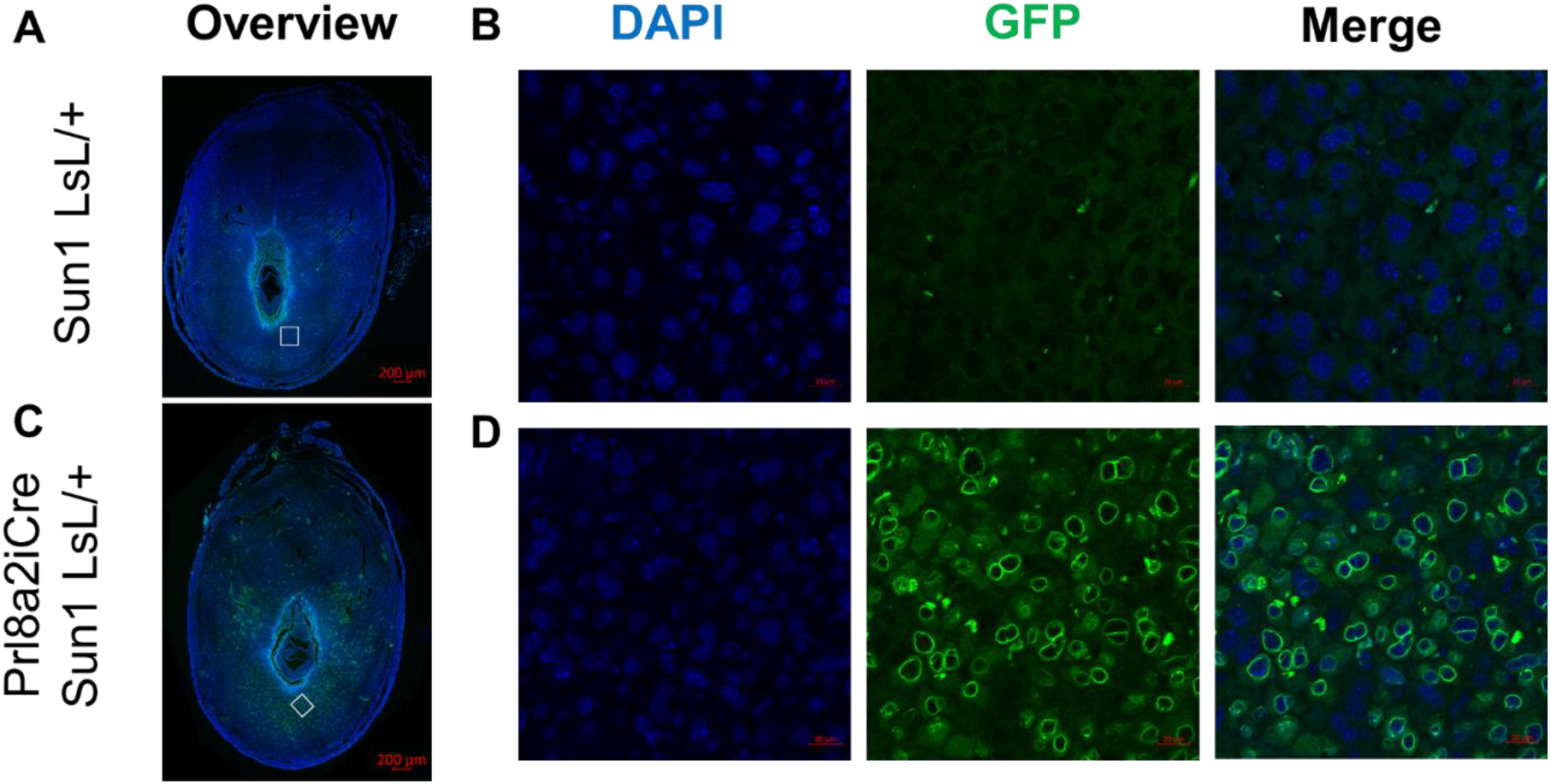
Uteri from gestational day 7.5 *Prl8a2*^*iC*re/+^ *Sun1*^LsL/+^ mice. *Prl8a2*^*i*Cre/+^ *Sun1*^LsL/+^ mice were mated with wild-type males and collected at gestational day (GD) 7.5 to validate iCre activity in decidual cells. A cross-section of an implantation site from *Sun1*^LsL/+^ mice (**A**) and *Prl8a2*^*i*Cre/+^ *Sun1*^LsL/+^ mice (**C**) were examined. The white box indicates the magnified section (**B**, **D**). Blue indicates DAPI nuclear stain and green indicates GFP stain. Scale bar represents 200 μm on overview image and 20 μm on magnified regions.

The spatial and temporal GFP expression pattern of *Prl8a2*^*iC*re/+^ driven GFP reporter is in line with that of endogenous *Prl8a2* transcripts. Fluorescent *in situ* hybridization data demonstrate that *Prl8a2* expression was not detected in the cross-sections of the uterus at GD 4 but was apparent on GD 4.5 adjacent to the embryo (**Figure 4A-B**). *Prl8a2* expression increased surrounding the embryo at GD 5.5, with more cells expressing *Prl8a2* in the anti-mesometrial compared to mesometrial region (**Figure 4C**). By GD 7.5, most decidua cells surrounding the embryo in the anti-mesometrial region express *Prl8a2* (**Figure 4D**). Our results, in agreeance with previous work (Alam et al., 2007), recapitulated the spatial expression pattern of the endogenous *Prl8a2* gene in the pregnant uterus. Of note, we detected *Prl8a2* expression at GD 4.5, which is earlier in gestation by a day compared to previous work (Alam et al., 2007).

**Figure 4.**
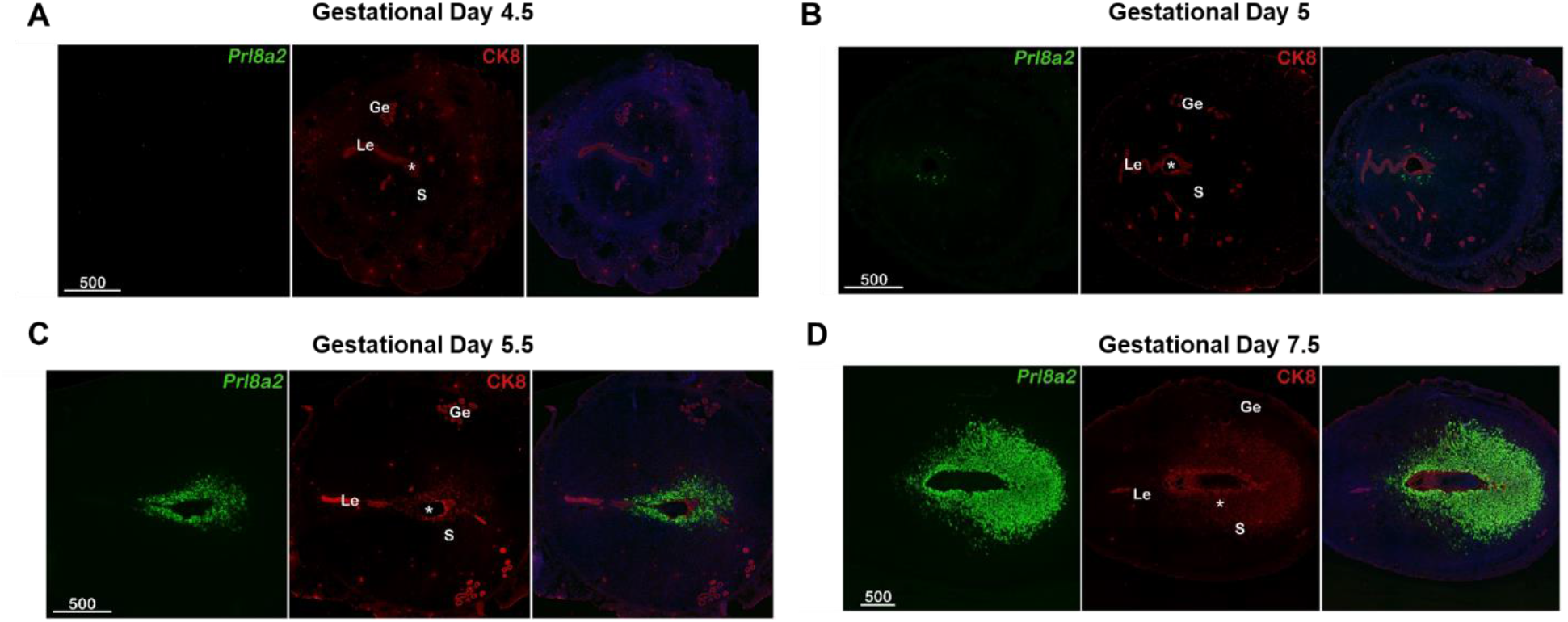
Fluorescent *in situ* hybridization for *Prl8a2* mRNA from cross-sectional mouse uteri. Mice were mated and tissues collected at gestational day 4.5 (**A**), 5 (**B**), 5.5 (**C**), and 7.5 (**D**). Fluorescent *in situ* hybridization depicts expression of *Prl8a2* (green) and cytokeratin (*CK*; red) in decidua over time. * indicates embryo and scale bar is 500 μm. LE = luminal epithelium, GE = Glandular epithelium, S = stroma.

Additionally, we wanted to ensure there were no off-target results from Prl8a2-iCre recombinase activity. Expression of *Prl8a2*^*iC*re/+^ driven GFP was also evaluated in the ovary, oviduct, pituitary and skeletal muscle. There was no GFP expression observed in ovary, oviduct, pituitary, or skeletal muscle (**Supplemental Figure 1**), supporting the tissue specificity of the Prl8a2-iCre in the decidua.

In line with the objective, we successfully created transgenic mice with iCre recombinase activity under the control of decidual cell marker, *Prl8a2*. This iCre is active during the post-implantation stages in the mouse, specifically at GD 7.5. In conclusion, the use of this mouse strain in future experiments could provide new insights into cellular mechanisms responsible for improper decidualization and implantation failure.

## Methods

All animal studies were conducted in accordance with the Guide for the Care and Use of Laboratory Animals published by the National Institutes of Health and animal protocols approved by the Animal Care and Use Committee (ACUC) at the National Institute of Environmental Health Sciences.

### Generation of transgenic mice

The targeting construct was generated on the PUC19 backbone with the iCre open reading frame and the polyA sequences from the pCAG-iCre plasmid (Addgene 89573) (Weinberg et al., 2017) and arms for homologous recombination from genomic DNA of mouse embryonic stem (ES) cells. The NEBuilder High-Fidelity DNA Assembly Cloning Kit (New England Biolabs, E5520) was used to assemble DNA fragments into the targeting construct based on manufacturer’s instructions with the primers detailed in **Supplemental Table 1**. Once assembled, the portion of the iCre open reading frame, the polyA sequence and two homologous arms was subject to Sanger sequencing to select for clones that have correct nucleotide sequences.

The *Prl8a2-iCre* locus was generated by replacing the genomic sequence immediately downstream of the start codon in exon 1 with the transgene (**Figure 1**). In the final plasmid donor, the 5’ homology arm corresponds to chr13:27,528,750-27,529,719 GRCm39 and the 3’ homology arm corresponds to chr13:27,531,356-27,532,420 GRCm39. The overall locus structure was maintained by replacing 1636 bp of endogenous sequence with the iCre-pA genetic payload. Gene targeting was done in B6129F1 embryonic stem cells (G4; 129S6/SvEvTac x C57BL/6Ncr). Embryonic stem cells were transfected with a 6:1 molar ratio of donor plasmid and Cas9-Puro/sgRNA(ACTAGCAGTCTTATCCAACC<u>CGG</u>) delivery plasmid (pSpCas9(BB)-2A-Puro (PX459) V2.0), a gift from Feng Zhang (Addgene plasmid # 62988, (Ran et al., 2013).

After transfection, the cells were exposed to 48-hours of puromycin selection (0.9 μg/mL) followed by standard clonal expansion/screening. Clones were screened with 5’ and 3’ screens external to the homology arms and genetic payload/WT zygosity screen to identify Prl8a2-iCre/WT clones. Screening primers: Prl8a2-Cre 5’Scr (Fwd: GTGGGACTATGACCCTATTGCTTAT, Rev: ATCTTCCAGGTGTGTTCAGAGA), Prl8a2-Cre 3’Scr (Fwd: CACTCGGAAGGACATATGGGAG, Rev: ACATTGAAGTTTAAGCTCATTTTCAG), and Prl8a2-Cre WT Scr (Fwd:, Rev:TGTGATGTCACTTGGCTTGTACT). Screening amplicons from targeted clones were fully sequenced to confirm proper insertion of the iCre-pA construct. Targeted ES cells were microinjected into albino B6J blastocyst for chimeric founder generation. The *Prl8a2-iCre* allele was re-sequenced in the F1 chimera offspring. The line was then crossed to C57BL/6J wildtype mice to establish and expand the colony. The final mouse line was genotyped at Transnetyx using primer/probe assays; Prl8a2 WT (Fwd: CCTCTCTCCCTGTTTTTGTTTGTTTTTTTAT, Rev: GAGGTATGGGATGTGAAACAATCA, Probe: CCCTCCTGGTCCACCC) and Prl8a2-iCre (Fwd: CAGAGTCTGAACTCATCCTGCTTGG, Rev: TCATCAGAGGTGGCATCCACAG, Probe: TCCTGGCCAATAGCC). These mice are available upon request and will be sent to the Jackson laboratory.

### Verification of iCre recombinase activity

The double fluorescent Cre recombinase reporter mouse, mTmG was established previously (Muzumdar et al., 2007). Founder mTmG mice were purchased from the Jackson Laboratory (Bar Harbor, ME). In these mice, the mTmG reporter transgene is driven by a strong and ubiquitous pCA promoter from the well-characterized *ROSA26* locus. The mTmG expresses membrane-targeted tdTomato (mT) prior to Cre-mediated excision and membrane-targeted green fluorescent protein (mG) following Cre excision. The mTmG mice were crossed with *Prl8a2*^iCre/+^ mice to generate *Prl8a2*^iCre/+^ mTmG mice.

*Prl8a2*^*i*Cre/+^ mice were crossed with CAG-Sun1sfGFP (*Sun1*^LsL/+^) mice (Mo et al., 2015) that were purchased from the Jackson Laboratory to generate *Prl8a2*^*i*Cre/+^*Sun1*^LsL/+^ mice. These mice contain a coding sequence for nuclear membrane Sad1 and UNC84 domain containing 1 (SUN1) fused at its C-terminus to two copies of superfolder GFP (sfGFP) followed by six copies of Myc inserted in the *Gt(ROSA)26Sor* locus. In these mice, there is a Cre-dependent removal of a floxed stop cassette which allows expression of *Sun1-sfGFP-myc* resulting in a SUN1-GFP fusion protein.

*Prl8a2*^iCre/+^ mTmG and *Prl8a2*^*i*Cre/+^*Sun1*^LsL/+^ female mice (2-4 months old) were housed with wild-type (CD1) stud males and monitored each morning. The morning a seminal plug was detected in the vagina was defined as GD 0.5. Female mice were collected at various GD and were euthanized by CO_2_ inhalation followed by cervical dislocation. Tissues were fixed in 4% paraformaldehyde in PBS for 24 h at 4°C for histology.

### Deciduoma induction

Adult female mice (> 6 weeks old) were ovariectomized and rested for two weeks. Artificial decidualization was induced by daily subcutaneous injections of 100 ng estradiol for three days followed by two days of rest. After resting, a combination of 1 mg progesterone and 6.7 ng estradiol was injected subcutaneously for three days. Six hours after last injection, the mice were anaesthetized with isoflurane and oxygen, and intraluminal uterine injection of 50 μl sesame oil was performed. Following surgery, the mice continued to receive daily subcutaneous injections of 1 mg progesterone and 6.7 ng estradiol for 5 days after the intraluminal injection procedure for induction of deciduoma.

### Fluorescence in situ hybridization

Fluorescence *in situ* hybridization was performed as previously described (Yuan et al., 2019). In brief, implantation sites from three individual animals in each experimental group were collected. Frozen sections (12 μm) from three implantation sites from different females in each group were mounted onto poly-L-lysine-coated slides and fixed in 4% paraformaldehyde in PBS. Following acetylation and permeabilization, slides were hybridized with the DIG-labeled *Prl8a2* probes at 55°C overnight. After hybridization, slides were then washed, quenched in H_2_O_2_ (3%), and blocked in blocking buffer (1%). Anti-Dig-peroxidase was applied onto hybridized slides and color was developed by Tyramide signal amplification (TSA) Fluorescein according to the manufacturer’s instructions (PerkinElmer). Cytokeratin 8 (Iowa hybridoma bank, 1:100 dilution) was used to outline epithelial cells. Images presented are representative of three independent experiments.

### Tissue processing, histology, and immunofluorescence staining

Following paraformaldehyde fixation, tissues from *Prl8a2*^*i*Cre/+^ mTmG mice were cryoprotected in 10% sucrose (6 h – overnight) and 20% sucrose (6 h – overnight) at 4°C, and then embedded in Tissue-Tek OCT compound, while tissues from *Prl8a2*^*i*Cre/+^*Sun1*^LsL/+^ mice were placed in 70% ethanol for at least 24 hours, followed by dehydration and paraffin embedding. For cryopreserved tissues, samples were sectioned at 10 μm thickness using a Leica cryostat and were mounted on glass slides. The slides were washed with PBS and mounted in Vectashield Hardset antifade mounting medium containing DAPI (H-1500, Vector Laboratories, Burlingame, CA). For paraffin embedded tissues, samples were sectioned at 5 μm thickness and mounted on glass slides. The slides were incubated in Citrosolv (Decon Labs Inc.) for three minutes thrice, followed by 100% ethanol three times, 95% and 70% once for one minute each. Antigen retrieval was performed by microwaving slides in a buffer containing Antigen Unmasking Solution (Vector H-3300) for 3.5 minutes at 70% power followed by 12 minutes at 20% power. Slides were cooled back to room temperature then washed with PBS once for 5 minutes, then PBS containing 0.2% triton-X for 10 min, then washed again with PBS for 5 minutes. Slides were blocked in PBS containing 5% normal donkey serum for 1 hour at room temperature. Incubation with a primary antibody for GFP (Abcam, ab5450) occurred overnight at 4°C in a humidified chamber. Slides were washed three times for 5 minutes each in PBS with 0.1% Tween-20 (PBST) and incubated in a secondary antibody (1:500, Donkey anti-Goat IgG (H+L) Cross-Adsorbed Secondary Antibody, Alexa Fluor 488, ThermoFisher A-11055) for 1 hour at room temperature. Slides were washed again as detailed above, then mounting media containing DAPI (Vectashield H-1200) was applied.

Fluorescent images of uteri from *Prl8a2*^*i*Cre/+^ mTmG mice were captured using Zeiss AX10 microscope equipped with a digital imaging system while fluorescent images of decidua from *Prl8a2*^*i*Cre/+^*Sun1*^LsL/+^ female mice were captured using ZEISS LSM 780 UV confocal microscope at 20X magnification.

## Acknowledgements

We would like to acknowledge Amanda Bartos, Xiaofei Sun, and Sudhansu Dey for their assistance with fluorescent in situ hybridization. We would also like to thank Olivia Emery for her assistance in the lab. This work was supported by R01HD096266 (T.E.S., A.M.K.), Korea Health Industry Development Institute (KHIDI), Grant/Award Number: HI17C2134 (Y.O.), and an Intramural Research Program of the NIEHS, United States, NIH project nos. Z1AES103311 (F.J.D.).

