## Supplemental Data for "Inserting Cre Recombinase into the Prolactin 8a2 gene for Decidua-Specific recombination in Mice"

Supplemental Figure 1.

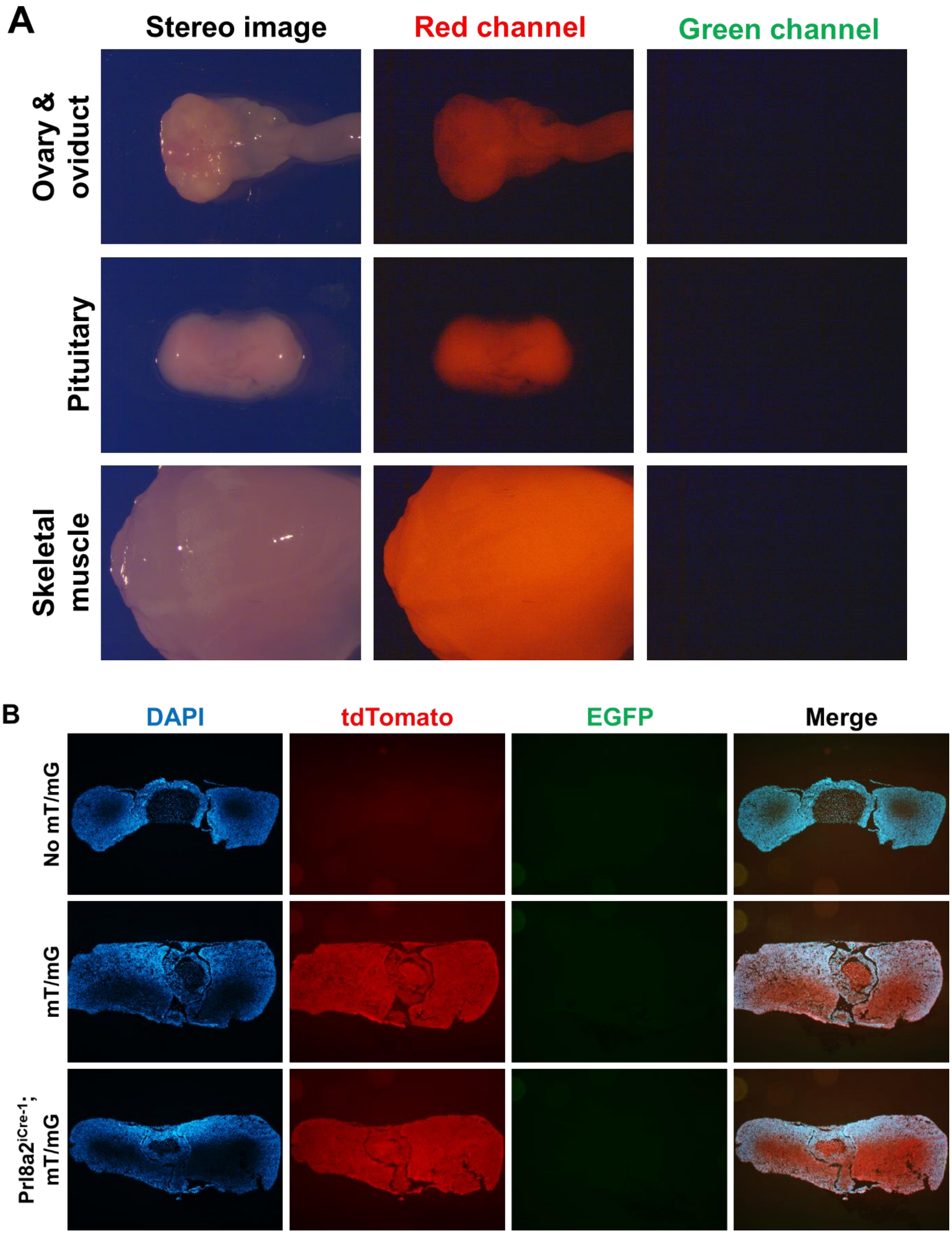

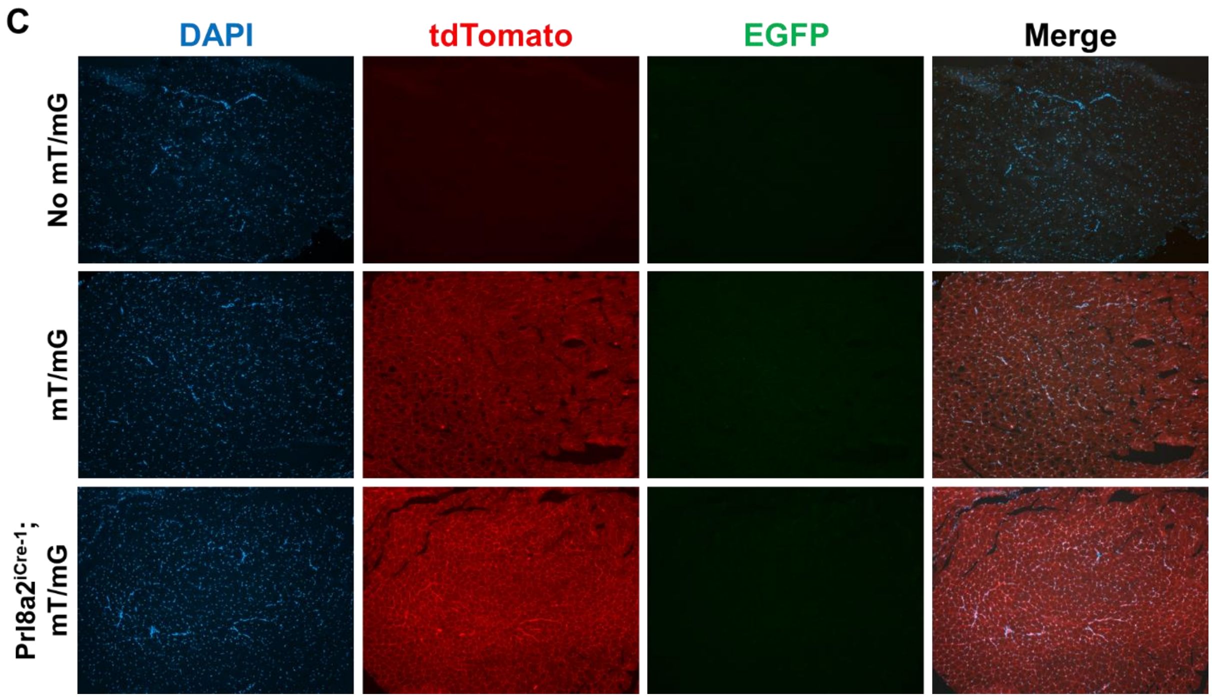


**Supplemental Figure 1. Images of pituitary, skeletal muscle, ovary and oviduct in *Prl8a2*$^{icre/+}$ mTmG mice.**

Various tissues were collected from *Prl8a2*$^{iCre/+}$ mTmG mice to confirm Prl8a2iCre specificity. There was no *Prl8a2*$^{iCre}$ activity visible in whole- mount observation in ovary, oviduct, pituitary, or skeletal muscle (**A**). Pituitary (**B**) and skeletal muscle (**C**) were sectioned and stained for EGFP. Blue indicates nuclear stain, DAPI.

**Supplemental Table 1. Primers used for targeting genetic construct.**

| Overlaps | Oligo | Anneals | Direction |
|---|---|---|---|
| pUC19 | tgcaggtcgactctagaggatccccTTGTACTGCCTAGTTTTGTGTCAACTTG | Left_arm | Forward |
| iCre_pA | ccatggtggcggcTATTGGCCAGGACTTTCCAAGCAGG | Left_arm | Reverse |
| Left_arm | gtcctggccaataGCCGCCACCATGGTGCCCAAGAAGA | iCre_pA | Forward |
| Right_arm | ggaactgcaacccTGCAGGTCGAGGGATCTCCATAAGAGAAGAGG | iCre_pA | Reverse |
| iCre_pA | ccctcgacctgcaGGGTTGCAGTTCCCTTTAGCTTCTTG | Right_arm | Forward |
| pUC19 | gccagtgaattcgagctcggtacccTGATGGGACAAGGACAGTGCAGCTT | Right_arm | Reverse |
